# Chaperone-Directed Ribosome Repair after Oxidative Damage

**DOI:** 10.1101/2022.07.28.501866

**Authors:** Yoon-Mo Yang, Youngeun Jung, Daniel Abegg, Alexander Adibekian, Kate Carroll, Katrin Karbstein

**Affiliations:** Department of Integrative Structural and Computational Biology, UF Scripps Biomedical Research, Jupiter, Florida 33458, United States of America; Department of Chemistry, UF Scripps Biomedical Research, Jupiter, Florida 33458, United States of America; HHMI Faculty Scholar, Chevy Chase, Maryland 20815, United States of America

## Abstract

Reactive oxygen species are ubiquitous in cells, where they damage RNA and protein. While relief mechanisms, including effects on translation, have been described, whether ribosomes are functionally compromised by oxidation, and how this damage is mitigated, remains unknown. Here we show that cysteines in ribosomal proteins, including Rps26, are readily oxidized and rendered non-functional, which is exacerbated when yeast are exposed to H_2_O_2_. Oxidized Rps26 is released from ribosomes by its chaperone Tsr2, which allows for repair of the damaged ribosomes with newly made Rps26. Ribosomes containing damaged Rpl10 or Rpl23 are similarly repaired by their chaperones, Sqt1 and Bcp1. Ablation of this pathway impairs growth, which is exacerbated under oxidative stress. These findings reveal a novel mechanism for chaperone-mediated ribosome repair with implications for aging and health.

**One-Sentence Summary:** Chaperones repair thiol-oxidized ribosomes by release of damaged components and incorporation of newly made ribosomal proteins.

## Main Text

Ribosomes are the RNA-protein complexes that synthesize proteins in all cells. Their functionality is carefully safeguarded, both during assembly (*1-4*) and their functional cycle (*5*), reflecting the importance of ribosomes in maintaining protein homeostasis. Equally important as maintaining correctly assembled and fully functional ribosomes is the need to maintain proper ribosome numbers, as changes lead to mRNA-specific changes in translation (*6*). As a result, cells devote more than half of all transcription and translation events to the synthesis of ribosomes (*7*), even though ribosomes are exceptionally stable (*5*).

Oxidative stress (OS) is both ubiquitous in actively growing cells, and exacerbated under specific conditions, which include the defense against intruders (for microorganisms), T-cell activation and increased autophagic flux (*8, 9*). All cellular biomolecules, including RNA and protein, are susceptible to oxidation, and due to their large size and abundance, ribosomes should be a major target of oxidation. How cells ameliorate damage from ribosome oxidation, while maintaining enough ribosomes, especially given that ribosome synthesis is down-regulated (*10*), is unknown. Notably, most oxidized proteins undergo degradation with few repairable proteins (*11*), and oxidized mRNAs are also decayed (*12*).

To test if ribosomal proteins (RPs) are damaged in yeast cells exposed to oxidative stress, we utilized a chemical probe for detecting cysteinyl oxidation in situ, called BTD (*13*), which tags sulfenic acid, the direct product of thiolate oxidation by H_2_O_2_. To investigate the link between OS and ribosome *S*-sulfenation we selected an initial concentration of H_2_O_2_ that activates a defined set of redox pathways (*14, 15*). Accordingly, live cells were treated with 1 mM H_2_O_2_, followed by a biotinylated BTD-analog (bio-BTD) for facile detection. Lysates were subsequently prepared and resolved on sucrose gradients to separate ribosomes. Western blotting demonstrates BTD-tagged proteins of the size expected for RPs in the 80S peak (**Fig. 1A**, top), and by blotting for individual RPs we identified Rps26 as a major BTD-reactive band (**Fig. 1A**, middle and bottom), indicating its susceptibility to oxidation. Of note, several bands, including that for Rps26, were labeled by BTD in the absence of exogenous peroxide, which is consistent with basal ribosome oxidation in growing cells. These findings demonstrate the susceptibility of Rps26 to cysteinyl oxidation, consistent with previous proteome-wide findings (*16-18*).

**Fig. 1.**
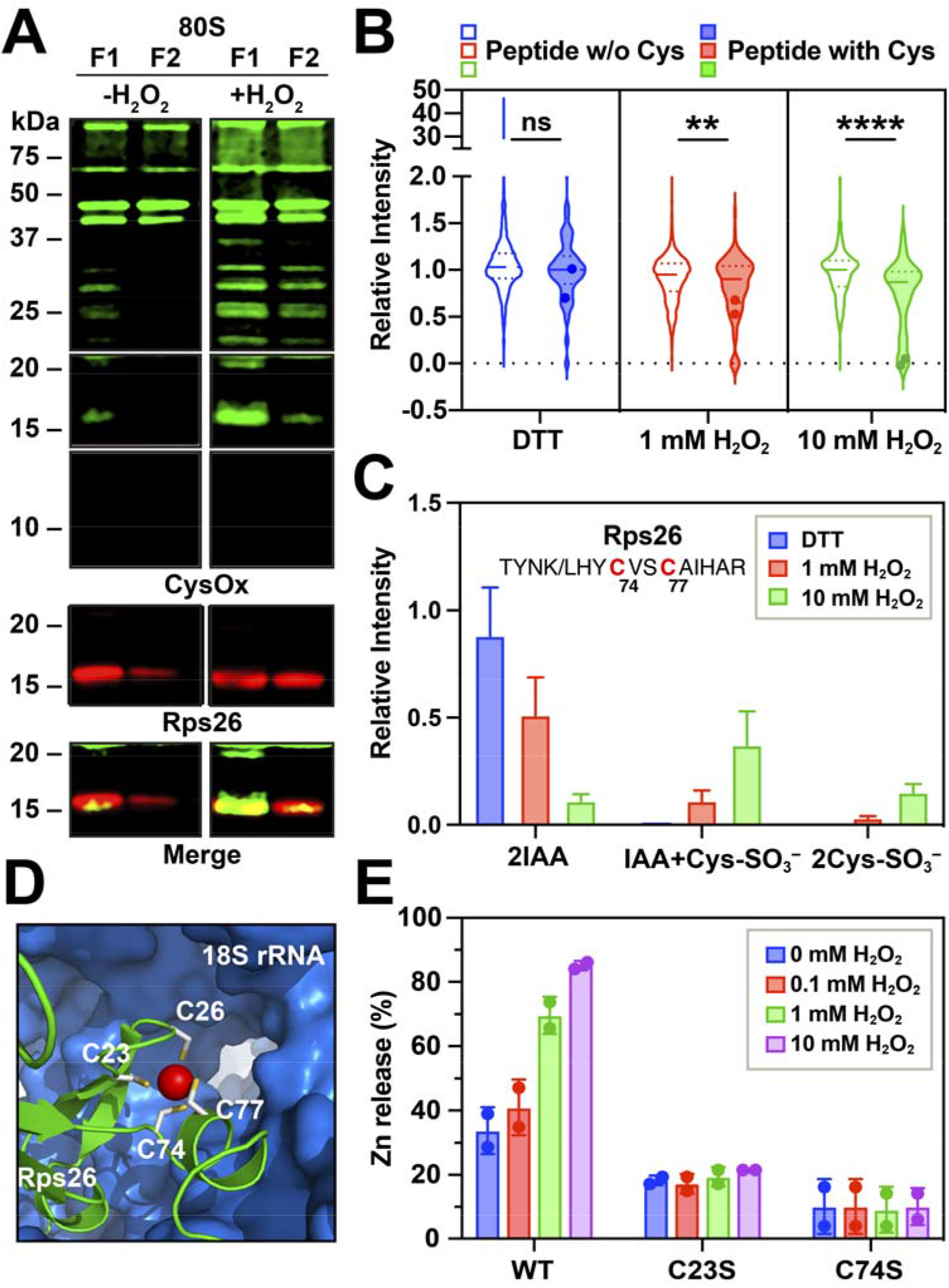
Oxidative stress damages Rps26. (A) BY4741 cells were treated with 1 mM H_2_O_2_ followed by 1 mM BTD treatment. The two fractions containing 80S ribosomes were collected from 10-50% sucrose gradients of cell lysates. (B) Peptides with and without cysteine residues derived from ribosomal proteins from purified 80S ribosomes. Rps26-derived peptides containing Cys74/Cys77 are indicated in dots. As in previous work (*16*), the other Cys-containing peptide was not detected. (C) Analysis of Rps26_Cys74/Cys77-containing peptides. (D) Structure of Rps26, indicating the Zn-finger (PDB 4V88). (E) H_2_O_2_-dependent Zn^2+^-release from Rps26. Release of Zn^2+^ from recombinant MBP-Rps26 variants was measured by monitoring formation of the Zn^2+^-PAR complex.

To independently confirm preferential oxidation of Rps26, and identify the sites of oxidation, we purified ribosomes, exposed them to H_2_O_2_, before labeling them with iodoacetamide (IAA). IAA reacts with thiolates and a decrease in alkylation, which can be identified in subsequent mass-spectrometry experiments, is indicative of cysteine oxidation. Among the peptides identified by LC-MS/MS, approximately ∼9% or 13% of cysteines showed a decrease in IAA-labeling after treatment with 1 mM or 10 mM H_2_O_2_ treatment, respectively (**Fig. 1B)**, indicating that most cysteines in assembled RPs are resistant to oxidation by H_2_O_2_. In contrast, 50% and 90% of cysteine-containing peptides in Rps26 had decreased IAA labeling at 1 and 10 mM H_2_O_2_, respectively (**Fig. 1C**). In particular, we identified Rps26 cysteines 74 and 77 as sites of oxidation. Furthermore, Rps26 was the only RP where decreased IAA labeling was accompanied by an increase in peptides containing a hyperoxidized form of cysteine (–SO_3_H). This proteomic analysis, together with *in situ* probe labeling, demonstrate that Rps26 is preferentially oxidized *in vivo* and *in vitro*.

Rps26 contains 4 cysteines, which chelate a Zn^2+^ ion in a structurally important Zn-finger motif (**Fig. 1D**). We therefore hypothesized that cysteinyl oxidation would lead to Zn^2+^ release, and inactivation of the protein. To test this hypothesis, we purified recombinant Rps26 (**Fig. S1A**), and quantified bound Zn and its release using the PAR-assay (*19*). Indeed, addition of increasing H_2_O_2_ concentrations increased the rate of Zn release, as well as the extent of Zn release **(Fig. 1E & S1D**), demonstrating Zn binding to ∼85% of the protein.

To test if the Zn-free protein is functional, we mutated each cysteine individually to serine, and tested for complementation of an Rps26-deficient yeast strain. This analysis showed that Cys23 and Cys74 are essential for cell viability (**Fig. S1B**). Moreover, purified recombinant Rps26_C23S and Rps26_C74S were essentially Zn-free (**Fig. 1E & S1C-E**). Cys26 was non-essential, and mutation of Cys77 led to a strong growth defect. Together, these data strongly suggest that Zn-binding to Rps26 is essential for its function.

Rps26 can be selectively released from ribosomes by the chaperone Tsr2 when its binding is weakened by loss of a Mg ion at the RNA-Rps26 interface, thereby producing Rps26-deficient ribosomes (*20*), which regulate metal ion homeostasis (*21*). Because the Zn-finger forms part of the 40S interface (**Fig. 1D**), we hypothesized that Tsr2 could similarly release oxidized Rps26 lacking Zn. To examine this idea, we incubated recombinant, purified Tsr2 with untreated 40S, or 40S exposed to 1 or 10 mM H_2_O_2_, and then assayed for Rps26 release (*20*). After oxidation, Rps26 was released from 40S subunits, and the extent of release increased with H_2_O_2_ concentration (**Fig. 2A**). Release was specific for Rps26 and not observed for Rps10, and required the ability of Tsr2 to bind both Rps26 (Tsr2_DWI, (*20, 22*)) and the 40S subunit (Tsr2_K/E, (*20*)), demonstrating active release, rather than ‘catching’ of released Rps26 (**Fig. 2B)**. Moreover, 40S oxidation, and not Tsr2 oxidation was required for this process (**Fig. S2A)**. Furthermore, H_2_O_2_- and chaperone-dependent release of Rps26 from the 40S subunit was specific to Tsr2/Rps26, as neither Rps2 nor Rps3 were released by their chaperones, Tsr4 and Yar1, respectively (**Fig. S2B-C**). Thus, these data demonstrate specific Tsr2-dependent release of oxidized Rps26 from fully assembled 40S ribosomes *in vitro*.

**Fig. 2.**
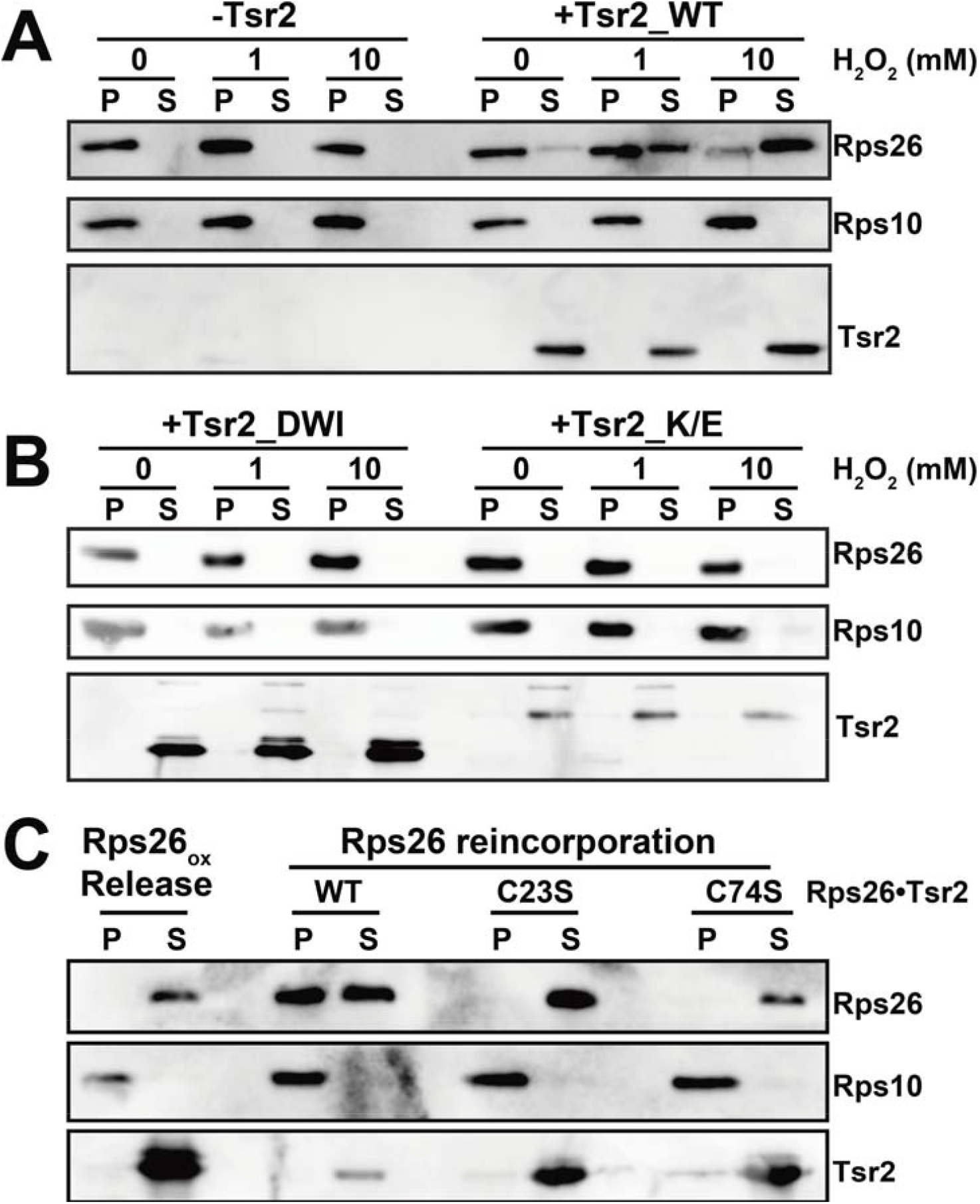
Tsr2 releases oxidized Rps26. (A) Western blot analysis of pelleted (P) ribosomes and proteins released into the supernatant (S). 200 nM purified 40S were incubated with or without 4 µM recombinant Tsr2 at the indicated H_2_O_2_ concentrations for 30 min, before assaying for Rps26 release. (B) Rps26 release assay with the Rps26-interaction-deficient Tsr2_DWI mutant or 40S-interaction-deficient Tsr2_K/E mutant. (C) Incorporation of the indicated Rps26 variants from Rps26•Tsr2 into ribosomes assayed by co-sedimentation with ribosomes.

We next investigated whether oxidized 40S could be repaired by replacing Rps26. For this, we incubated ribosomes from which damaged Rps26 was released with recombinant Rps26•Tsr2. Tsr2 was able to deliver Rps26 to 40S, showing that oxidized Rps26 in damaged ribosomes can be replaced with newly made Rps26. By contrast, Zn-less Rps26_C23S and Rps26_C74S cannot associate with these ribosomes (**Fig. 2C**). Thus, Tsr2 extracts non-functional Zn-less Rps26 and replaces it with intact Zn-bound Rps26.

To examine this process *in vivo*, we tested whether Tsr2 binding to ribosomes increased with H_2_O_2_ treatment. Indeed, sucrose gradient fractionation demonstrates that exposure of yeast to H_2_O_2_ leads to increased Tsr2 binding to 80S ribosomes (**Fig. 3A**). This finding is specific to Tsr2, and not observed for the Rps3 chaperone Yar1, and complemented by *in vitro* binding studies showing increased affinity of Tsr2 but not Yar1 to oxidized 40S (**Fig. S2E-F**).

**Fig. 3.**
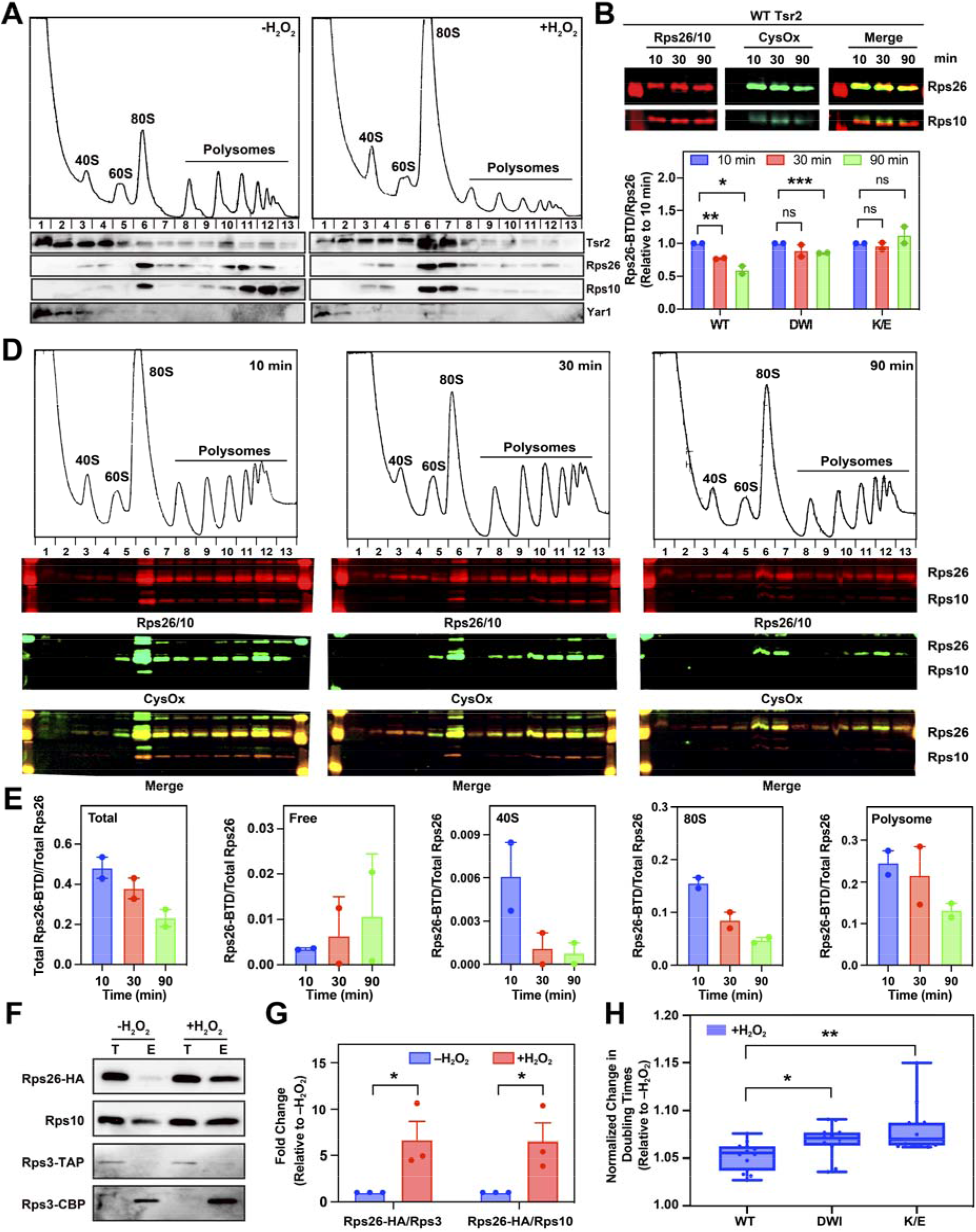
Oxidized Rps26 is released from ribosomes before being degraded. (A) Tsr2 binding to ribosomes under oxidative stress. 10-50% sucrose gradients of cell lysates from BY4741 cells with (right) or without (left) 1 mM H_2_O_2_ treatment for 30 min. Absorbance profile at 254 nm (top) with Western blots (bottom). (B) Western blots of purified ribosomes after pulse-chase BTD labeling shows decay of oxidized Rps26. (C) Quantification of BTD-labeled Rps26 in panel B and S3A. Data are the average of two biological replicates, and error bars indicate SEM. *p < 0.05 by unpaired t-test. (D) 10-50% sucrose gradients of pulse-chase BTD-labeled cells. (E) Quantification of BTD-labeled Rps26 in panel (D). Data are the average of two biological replicates, and error bars indicate SEM. (F) Newly-made Rps26-HA is incorporated into old ribosomes (Rps3-TAP, **Fig. S3E**). Pre-existing ribosomes were isolated using Rps3-TAP affinity purification (*20, 21*) and incorporation of newly-made Rps26-HA was measured by Western blot with or without treatment with 1 mM H_2_O_2_. T: total lysate; E: elution. TEV-protease elution from the IgG beads converts Rps3-TAP (Rps3-CBP-proteinA) to Rps3-CBP (Rps3-calmodulin-binding protein). (G) Quantification of three replicates of data in panel (F). (H) Changes in doubling time of cells expressing the indicated Tsr2 variants after exposure to 1 mM H_2_O_2_ in glucose media, relative to growth in untreated conditions. Data are the average of four biological replicates with three technical replicates each, and error bars indicate SEM. *p < 0.05, **p < 0.01 by unpaired t-test.

To test if the oxidized Rps26 is released from 40S subunits *in vivo*, we utilized the bio-BTD to label the oxidized cysteine (*13*). If Rps26 is released after exposure to H_2_O_2_, we would expect the oxidized Rps26 to disappear over time. Indeed, the amount of oxidized Rps26 relative to total Rps26 decreased over time. Importantly, this occurs in a Tsr2-dependent manner, as the amount of oxidized Rps26 remained unchanged in the non-functional Tsr2 mutants (DWI or K/E, **Fig. 3B-C & S3A**), strongly suggesting that the decay of oxidized Rps26 reflects degradation of Rps26 and not the entire ribosome and demonstrating that Tsr2 is required for Rps26 turnover.

To independently verify that Rps26 is first released from ribosomes before being degraded, we used a pulse-chase experiment to follow the ribosome binding of oxidized Rps26 over time. After exposure to exogenous peroxide, cells were incubated with bio-BTD, collected by centrifugation and resuspended in rich media without H_2_O_2_ and bio-BTD for 10, 30 or 90 minutes. Cells were then harvested, lysates generated and fractionated on sucrose gradients (**Fig. 3D)**. Analyzing the distribution of oxidized Rps26 over time demonstrates that its disappearance from 40S and 80S ribosomes is faster than its turnover from cells as a whole (**Fig. 3E)**. Moreover, the decrease of oxidized Rps26 in 40S and 80S subunits is accompanied by an initial increase in free oxidized Rps26. Thus, oxidized Rps26 is first released from ribosomes, before it is degraded.

Next, we wanted to test if release of damaged Rps26 allowed for repair of the ribosomes with newly made Rps26. To investigate this possibility we carried out a different pulse-chase experiment (**Fig. 3F & S3E**), where exposure to H_2_O_2_ was coupled to a switch from expression of untagged Rps26 to HA-tagged Rps26, which is fully functional (*20*) and can be readily distinguished by Western blotting. Moreover, ribosomes made prior to H_2_O_2_ addition were traced with Rps3-TAP, which allows for affinity purification (*21*). Thus, by purifying old ribosomes using the TAP-tag, and probing for the presence of Rps26-HA, we could then ask if new Rps26 was incorporated into old ribosomes when cells were exposed to oxidative stress. Indeed, >6-fold more newly-made Rps26-HA incorporated into pre-made ribosomes after H_2_O_2_ treatment (**Fig. 3F-G**). Thus, these data demonstrate that damaged ribosomes containing oxidized Rps26 are repaired with newly made, functional Rps26 *in vivo*. Importantly, these data also show that some Rps26-repair occurs even without H_2_O_2_ treatment, indicating the importance of this mechanism for maintenance of functional 40S ribosomes.

To examine the importance of ribosome repair of Rps26-damaged ribosomes, we next tested whether inactivation of Tsr2 increases sensitivity to H_2_O_2_. Indeed, yeast expressing the inactive Tsr2 mutants (DWI or K/E) are more sensitive to H_2_O_2_, even when they are overexpressing Rps26 to maintain full Rps26 occupancy (**Fig. 3H & S3C, F**). Moreover, repair of the Rps26-free ribosomes is required for the oxidative stress resistance as Rps26-deficient ribosomes are detrimental to the oxidative stress response (**Fig. S3G-H**). Finally, in the absence of H_2_O_2_, Tsr2 mutant cells grow more slowly, even when Rps26 is overexpressed to compensate for effects on Rps26 incorporation (**Figure S3B**, (*20, 22*)), again supporting the notion that Rps26 damage and the requirement for its repair are constitutive in growing cells.

To test the generality of this model, we next asked if other damaged RPs can be similarly released for ribosome repair. We expressed and purified several other chaperones (*23-27*) and incubated them with either the 40S or 60S subunit in the presence of H_2_O_2_. These studies show that oxidized Rpl10 and Rpl23 can be released by their cognate chaperones, Sqt1 and Bcp1, respectively (**Fig. 4A-B**), while Tsr4, Yar1 or Puf6/Loc1 did not release their RPs (**Fig. S2B-C & S4A)**. Interestingly, significant oxidation of Rpl10 was observed by MS (**Fig. S4B**), as previously observed (*16-18*). In contrast to previous work (*16*), we were unable to observe significant oxidation of Rpl23 (**Fig. S4C**), reflecting the reduced release of Rpl23 relative to Rpl10. Finally, pre-treatment of 60S but not Bcp1 with H_2_O_2_ leads to release of Rpl23, consistent with oxidation of Rpl23 and not Bcp1 (**Fig. S4D**), and increased interaction of Bcp1 with purified ribosomes was observed in the presence of H_2_O_2_ (**Fig. S4E**).

**Fig. 4.**
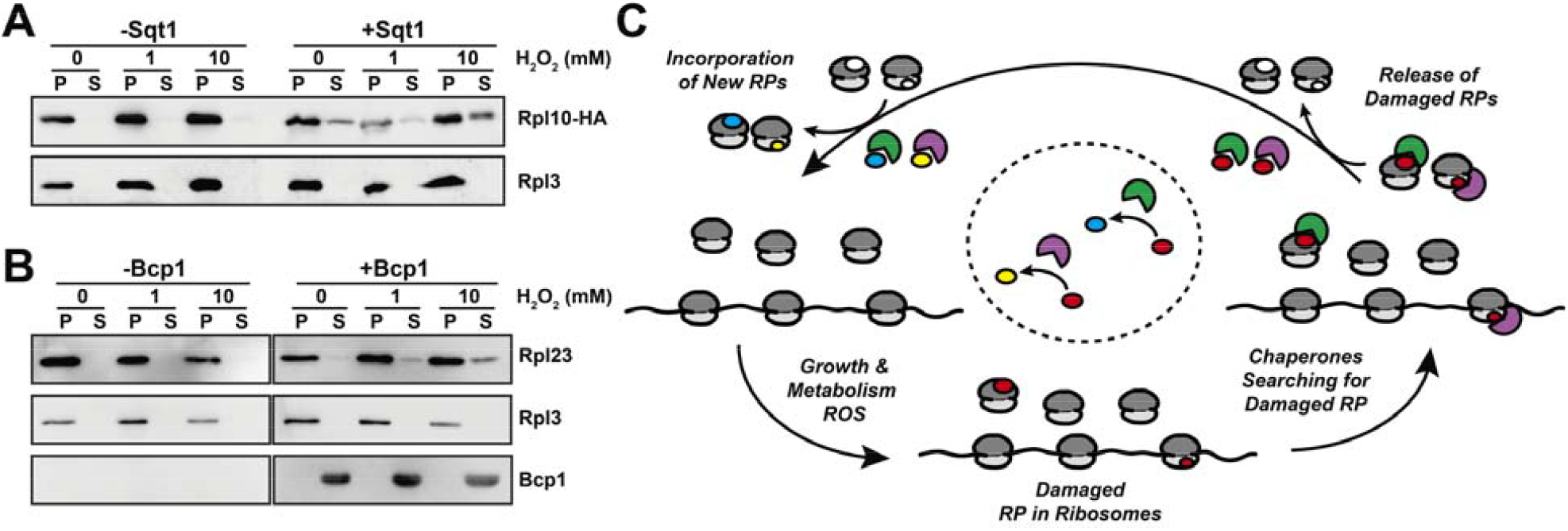
Chaperone-mediated release of 60S proteins. (A-B) 200 nM mature 60S subunits were incubated with or without 4 μM recombinant Sqt1 (A) or Bcp1 (B) and release of Rpl10 (A) or Rpl23 (B) from the ribosome pellet (P) into the supernatant (S) were tested using the pelleting release assay. (C) Model of chaperone-directed ribosome repair.

In summary, our findings demonstrate the preferred oxidation of Rps26, Rpl10, and to a lesser extent Rpl23, revealing the existence of a previously unrecognized ribosome repair pathway, in which individual chaperones direct the replacement of cognate oxidized RPs (**Fig. 4C**), thereby rationalizing their existence. The data support a mechanism where chaperones release their cognate RPs when their binding is compromised due to oxidative damage. Newly made RPs are then incorporated to repair the subunit. Importantly, the ability to bind the ribosome is required for this activity, showing that chaperones do not just simply replace already released RPs. Inactivating the repair machinery of just one RP, Rps26, leads to growth defects that are exacerbated under oxidative stress, demonstrating the importance of this pathway for ribosome maintenance.

## Acknowledgments

We thank V. Panse, M. Seedorf, J. Warner, B. Pertschy and KY Lo for generous gifts of antibodies.

## Author contributions

Y.Y. carried out the experiments.

Y.J. and K.C. prepared biotinylated BDT-probes.

D.A. and A.A. carried out the Mass spectrometry experiment and analysis.

Y.Y. and K.K. designed and analyzed the experiments and wrote the paper.

## Data and materials availability

All data are available in the main text or the supplementary materials.

## Supplementary Materials

Materials and Methods

Figs. S1 to S4

Tables S1 to S2

Data S1

References (*28*–*35*)

## References and Notes

1. H. Huang, H. Ghalei, K. Karbstein, Quality control of 40S ribosome head assembly ensures scanning competence. J Cell Biol 219, (2020).

2. M. D. Parker, J. C. Collins, B. Korona, H. Ghalei, K. Karbstein, A kinase-dependent checkpoint prevents escape of immature ribosomes into the translating pool. PLoS Biol 17, e3000329 (2019).

3. B. S. Strunk, M. N. Novak, C. L. Young, K. Karbstein, A translation-like cycle is a quality control checkpoint for maturing 40S ribosome subunits. Cell 150, 111–121 (2012).

4. H. Ghalei et al., The ATPase Fap7 Tests the Ability to Carry Out Translocation-like Conformational Changes and Releases Dim1 during 40S Ribosome Maturation. Mol Cell 68, 1155 (2017).

5. F. J. LaRiviere, S. E. Cole, D. J. Ferullo, M. J. Moore, A late-acting quality control process for mature eukaryotic rRNAs. Mol Cell 24, 619–626 (2006).

6. E. W. Mills, R. Green, Ribosomopathies: There’s strength in numbers. Science 358, (2017).

7. J. R. Warner, The economics of ribosome biosynthesis in yeast. Trends Biochem Sci 24, 437–440 (1999).

8. R. Scherz-Shouval, Z. Elazar, Regulation of autophagy by ROS: physiology and pathology. Trends Biochem Sci 36, 30–38 (2011).

9. D. G. Franchina, C. Dostert, D. Brenner, Reactive Oxygen Species: Involvement in T Cell Signaling and Metabolism. Trends Immunol 39, 489–502 (2018).

10. A. P. Gasch et al., Genomic expression programs in the response of yeast cells to environmental changes. Mol Biol Cell 11, 4241–4257 (2000).

11. D. Reichmann, W. Voth, U. Jakob, Maintaining a Healthy Proteome during Oxidative Stress. Mol Cell 69, 203–213 (2018).

12. R. Parker, RNA degradation in Saccharomyces cerevisae. Genetics 191, 671–702 (2012).

13. V. Gupta, J. Yang, D. C. Liebler, K. S. Carroll, Diverse Redoxome Reactivity Profiles of Carbon Nucleophiles. J Am Chem Soc 139, 5588–5595 (2017).

14. A. Delaunay, D. Pflieger, M. B. Barrault, J. Vinh, M. B. Toledano, A thiol peroxidase is an H2O2 receptor and redox-transducer in gene activation. Cell 111, 471–481 (2002).

15. B. D’Autreaux, M. B. Toledano, ROS as signalling molecules: mechanisms that generate specificity in ROS homeostasis. Nat Rev Mol Cell Biol 8, 813–824 (2007).

16. U. Topf et al., Quantitative proteomics identifies redox switches for global translation modulation by mitochondrially produced reactive oxygen species. Nat Commun 9, 324 (2018).

17. J. Meng et al., Global profiling of distinct cysteine redox forms reveals wide-ranging redox regulation in C. elegans. Nat Commun 12, 1415 (2021).

18. S. Akter et al., Chemical proteomics reveals new targets of cysteine sulfinic acid reductase. Nat Chem Biol 14, 995–1004 (2018).

19. J. B. Hunt, S. H. Neece, A. Ginsburg, The use of 4-(2-pyridylazo)resorcinol in studies of zinc release from Escherichia coli aspartate transcarbamoylase. Anal Biochem 146, 150–157 (1985).

20. Y. M. Yang, K. Karbstein, The chaperone Tsr2 regulates Rps26 release and reincorporation from mature ribosomes to enable a reversible, ribosome-mediated response to stress. Sci Adv 8, eabl4386 (2022).

21. M. B. Ferretti, H. Ghalei, E. A. Ward, E. L. Potts, K. Karbstein, Rps26 directs mRNA-specific translation by recognition of Kozak sequence elements. Nat Struct Mol Biol 24, 700–707 (2017).

22. S. Schutz et al., Molecular basis for disassembly of an importin:ribosomal protein complex by the escortin Tsr2. Nat Commun 9, 3669 (2018).

23. M. West, J. B. Hedges, A. Chen, A. W. Johnson, Defining the order in which Nmd3p and Rpl10p load onto nascent 60S ribosomal subunits. Mol Cell Biol 25, 3802–3813 (2005).

24. I. Rossler et al., Tsr4 and Nap1, two novel members of the ribosomal protein chaperOME. Nucleic Acids Res 47, 6984–7002 (2019).

25. Y. T. Yang, Y. H. Ting, K. J. Liang, K. Y. Lo, The Roles of Puf6 and Loc1 in 60S Biogenesis Are Interdependent, and Both Are Required for Efficient Accommodation of Rpl43. J Biol Chem 291, 19312–19323 (2016).

26. B. Koch et al., Yar1 protects the ribosomal protein Rps3 from aggregation. J Biol Chem 287, 21806–21815 (2012).

27. Y. H. Ting et al., Bcp1 Is the Nuclear Chaperone of Rpl23 in Saccharomyces cerevisiae. J Biol Chem 292, 585–596 (2017).

28. M. S. Longtine et al., Additional modules for versatile and economical PCR-based gene deletion and modification in Saccharomyces cerevisiae. Yeast 14, 953–961 (1998).

29. J. W. Lee, J. D. Helmann, Biochemical characterization of the structural Zn2+ site in the Bacillus subtilis peroxide sensor PerR. J Biol Chem 281, 23567–23578 (2006).

30. M. G. Acker, S. E. Kolitz, S. F. Mitchell, J. S. Nanda, J. R. Lorsch, Reconstitution of yeast translation initiation. Methods Enzymol 430, 111–145 (2007).

31. J. C. Collins et al., Ribosome biogenesis factor Ltv1 chaperones the assembly of the small subunit head. J Cell Biol 217, 4141–4154 (2018).

32. A. D. Brunner et al., Ultra-high sensitivity mass spectrometry quantifies single-cell proteome changes upon perturbation. Mol Syst Biol 18, e10798 (2022).

33. J. Cox et al., Accurate proteome-wide label-free quantification by delayed normalization and maximal peptide ratio extraction, termed MaxLFQ. Mol Cell Proteomics 13, 2513–2526 (2014).

34. S. Schutz et al., A RanGTP-independent mechanism allows ribosomal protein nuclear import for ribosome assembly. Elife 3, e03473 (2014).

35. G. Poll et al., rRNA maturation in yeast cells depleted of large ribosomal subunit proteins. PLoS One 4, e8249 (2009).

