## Supplemental materials for "Chaperone-Directed Ribosome Repair after Oxidative Damage"

**This PDF file includes:**

Materials and Methods
Figs. S1 to S4
Tables S1 to S2
Data S1

### Materials and Methods

#### Strains and plasmids

*S. cerevisiae* strains used in this study were either purchased from the GE Dharmacon Yeast Knockout Collection or constructed using standard methods (*28*) and are listed in Table S1. The Gal::Rpl10 strain was a generous gift from Philipp Milkereit. Plasmids are listed in Table S2.

#### Protein purification

Rps26, Tsr2, Tsr2_DWI, Tsr2_K/E, Tsr4, Yar1 and Rps26·Tsr2 complex were purified as previously described (*20*). Bcp1, Sqt1, Puf6 and Loc1 were expressed in *E. coli* Rosetta2 (DE3) cells (Novagen) as TEV-cleavable His6-MBP fusion proteins. Cells were grown at 37 °C in LB medium supplemented with ampicillin. At $OD_{600}$ 0.4 protein expression was induced with 1 mM IPTG for 16 hours at 18 °C. Proteins were purified using Ni-NTA resin (Qiagen) according to the manufacturer's instructions. Eluted proteins were pooled and dialyzed overnight at 4 ˚C into 50 mM Tris (pH 7.4), 100 mM NaCl, and 1 mM DTT with tobacco etch virus (TEV) protease. Cleaved tag-less proteins were further purified by MonoQ/MonoS and Superdex 75 (GE) chromatography, in 50 mM Tris (pH 7.4), 100 mM NaCl and a linear gradient to 1M NaCl. For purification of the MBP-tagged Rps26 variants (wt or C23S or C74S), amylose resin (NEB) was used according to the manufacturer's instructions. Remaining maltose in elution buffer was removed during concentration into storage buffer (50 mM Tris (pH 7.4), 100 mM NaCl, 5% glycerol). Protein concentration was determined using absorption at 280 nm or via Bradford (for Rps26 and its variants), and proteins were stored at -80°C.

#### Zn release assay

$Zn^{2+}$ release was measured as described previously (*29*) using 100 µl of 3.2 µM MBP-tagged Rps26 variants in the presence of 100 µM 4-(2-pyridylazo)resorcinol (PAR) and the indicated $H_2O_2$ concentrations. $Zn^{2+}$-PAR complex was monitored by measuring absorbance at 494 nm every 5 seconds for 30 minutes in 96-well plates (Thermo Scientific) using a Synergy.2 plate reader (BioTek). The $Zn^{2+}$ content of purified MBP-tagged Rps26 variants was calculated using a molar extinction coefficient of 14,900 $M^{-1}$ $cm^{-1}$ for the $Zn^{2+}$-PAR complex.

#### Lysis of yeast cells

All yeast cells were harvested, washed, and then resuspended in 1 ml/g cell pellet of the appropriate lysis buffer. The suspension was frozen by dripping into liquid $N_2$ produce pearls, that were then ground with mortar and pestle under liquid $N_2$. The resulting powder was stored at -80°C until use. At that time, another 1 ml/g cell pellet of the appropriate lysis buffer was added together with ~ 0.5 g glass beads, and the mixture thawed while rotating. Cell debris and beads were removed by centrifugation for 10 minutes at 3000g, and the supernatant was clarified by centrifugation for 10 minutes at 20,000g.

#### Ribosome Purification

Ribosome purification was carried out essentially as previously described (*30, 31*). BY4741 cells were cultured in YPD and harvested at $OD_{600}$ ~1.8. The cell pellet was further washed and resuspended in 1 ml/g cell pellet of Ribosome buffer (20 mM Hepes/KOH (pH 7.4), 100 mM KOAc, 2.5 mM $Mg(OAc)_2$) supplemented with 1 mg/ml heparin, 1 mM benzamidine, 1 mM PMSF and complete protease inhibitor cocktail (Roche), and processed as described above.

3 ml of clarified lysate was layered over 500 µl of sucrose cushion (Ribosome Buffer, 500 mM KCl, 1 M sucrose, 2 mM DTT) and spun in a Beckman TLA 110 rotor at 70,000 rpm for 65 min. The resulting pellet was resuspended in high salt buffer (Ribosome Buffer, 500 mM KCl, 1 mg/ml heparin, 2 mM DTT) and layered over 500 µl of sucrose cushion and spun in a Beckman TLA 110 rotor at 100,000 rpm for 70 min. The pellet was resuspended in ribosome storage buffer (Ribosome Buffer, 250 mM sucrose) and stored at -80°.

To isolate individual subunits, the ribosome pellet after the second centrifugation was resuspended in subunit separation buffer (50 mM HEPES/KOH, pH 7.4, 500 mM KCl, 2 mM $MgCl_2$, and 2 mM DTT), and 1 mM puromycin (Sigma-Aldrich) was added, and incubated for 10 min at 37°C . Subunits were isolated by loading onto 5–20% sucrose gradients (50 mM HEPES/KOH, pH 7.4, 500 mM KCl, 5 mM $MgCl_2$, 2 mM DTT, and 0.1 mM EDTA) and centrifuged at 19,600 rpm for 16 h. 40S or 60S subunits were collected separately and buffer was exchanged into ribosome storage buffer during concentration with Amicon concentrators (100 kDa MWCO). Concentrations were calculated using an extinction coefficient 2 X $10^7$ and 4 X $10^7$ $M^{-1}$ $cm^{-1}$ at $OD_{260}$ for 40S and 60S subunits, respectively.

To detect Rpl10 release, ribosomes were purified yeast strain YKK1545 (Rpl10 k/o Gal::Rpl10) containing pKK30938 (TEF-Rpl10-HA).

***In vitr*o Rps26 release assay**

Release assays were performed by pelleting as previously described (*20*). Briefly, 4 µM purified recombinant chaperone (Tsr2, Tsr2_DWI, Tsr2_K/E, Tsr4, Yar1, Bcp1, Sqt1 or Puf6/Loc1) were mixed with 200 nM 40S or 60S subunits, incubated for 30 min at RT and 10 min on ice in binding/release buffer (20 mM HEPES, pH 7.3, 2.5 mM MgOAc, 500 mM KOAc, 0.1 mg/ml heparin and 0.5 µl RNasin [NEB]) containing the indicated concentrations of $H_2O_2$ (Sigma). Samples were layered onto a 400 µl sucrose cushion (ribosome binding/release buffer + 20% sucrose (w/v)) and spun for 2.5 hours at 400,000 × g at 4 °C in a TLA100.1 rotor (Beckman). Supernatants were precipitated using trichloroacetic acid and resuspended in the same volume as pellets. Total resuspended sample was loaded on SDS-PAGE followed by Western blotting.

***In vitro* ribosome binding of chaperones**

1 µM purified ribosomes were incubated for 30 minutes at room temperature with 40 µM recombinant chaperone in gradient buffer (20 mM HEPES, pH 7.4, 5 mM $MgCl_2$ and 100 mM KCl) containing 1 mM of $H_2O_2$ in 100 µl. Each sample was applied to 10-50% sucrose gradients, centrifuged in a SW41Ti rotor for 2hours at 40,000 rpm and then fractionated. Proteins were detected using Western blotting.

**Polysome profiling**

BY4741 cells were grown in YPD media to mid-log phase. At mid-log phase cells were further cultured with or without 1 mM of $H_2O_2$. After 30 minutes 0.1 mg/mL cycloheximide was directly added to the culture and cells were harvested by centrifugation. Cell pellet was further washed and lysed in gradient buffer (20 mM Hepes, pH 7.4, 5 mM $MgCl_2$, 100 mM KCl, and 2 mM DTT) supplemented with 0.1 mg/mL cycloheximide, 1 mM benzamidine, 1 mM PMSF and complete protease inhibitor cocktail (Roche). Cleared lysate was applied to 10-50% sucrose gradients and centrifuged in an SW41Ti rotor for 2h at 40,000 rpm and then fractionated. Western blot was performed to probe proteins as indicated.

**BTD-labeling**

Biotinylated BTD is a benzothiazine-derived probe, which covalently binds to the sulfenic acid form of oxidized cysteine (*13*), but is also biotinylated for Western blot detection. For steady state BTD labeling, BY4741 cells were grown in 1 L of YPD to mid-log phase and treated with 1 mM of $H_2O_2$ for 10 min before harvesting in the presence of 0.1 mg/mL cycloheximide. The cell pellet was further washed and incubated at RT for 5 min in gradient buffer supplemented with 1 mM BDT, 0.1 mg/mL cycloheximide, 1 mM benzamidine, 1 mM PMSF and complete protease inhibitor cocktail (Roche). Further gradient centrifugation with cell lysate was performed as described above.

For pulse-chase BDT labeling, BY4741 cells grown in 1 L of YPD were harvested at mid-log phase, resuspended in 1 ml fresh YPD containing 1 mM of BTD with or without 1 mM of $H_2O_2$ for 5 minutes. Cells were pelleted, washed and resuspended in 1 L of fresh YPD. At the indicated time points 0.1 mg/mL cycloheximide was directly added to culture and gradient centrifugation was performed as described above.

**Isolation of Rps3 TAP-tagged pre-made ribosomes**

1 L of YKK491 (Gal:Rps26A, *Δ*Rps26B) cells containing plasmids pKK31045 (Gal::Rps3-TAP) and pKK30999 ($TET_{off}$::Rps26-HA) were grown to mid-log phase in galactose media containing 200 ng/ml doxycycline and then shifted to glucose media for 1 hour. Next, cells were either treated with or without 1 mM $H_2O_2$ for 2 hours before harvest. After harvesting and lysis in IgG binding buffer (50mM Tris pH 7.5. 100 mM NaCl, 10 mM $MgCl_2$, 0.075% NP40, 1 mM benzamidine and 1 mM PMSF), ~200 µl of pre-equilibrated IgG sepharose bead slurry (GE) was added to each lysate and incubated for ~2 hours at 4 ºC. After binding, each sample was washed 3 times with IgG binding buffer. Elution was performed by incubation for ~2 hours at 16 ºC with TEV protease (1:100, Invitrogen), 0.5 mM EDTA and 1 mM DTT in 200 µl IgG binding buffer. Samples were further analyzed using Western blot.

#### Serial dilution

Cells were grown in appropriate glucose minimal media overnight, and then diluted into fresh YPD for ~2 hours. Cells were diluted to $OD_{600}$ 0.5 before spotted on glucose or galactose plates with 10-fold serial dilutions.

#### Quantitative yeast growth measurements

Gal::Tsr2 cells (YKK1109) supplemented with plasmids encoding the Tsr2 variants and excess Rps26 were grown in appropriate glucose minimal media overnight, and then diluted into fresh YPD for ~2 hours before inoculating into 96-well plates (Thermo Scientific) at a starting $OD_{600}$ between 0.04 and 0.1. The additional Rps26 was necessary because cells with Tsr2 mutants otherwise contain ribosomes lacking Rps26, which cause a growth defect (**Fig. S3B**). A Synergy.2 plate reader (BioTek) was used to record the $OD_{600}$ for 24 hours, while shaking at 30 ºC. Doubling times were calculated using data points within the mid-log phase using GraphPad Prism 9. Statistical analyses for each measurement are detailed in the respective figure legend.

#### Mass spectrometry

500 µl of 200 nM purified ribosomes were incubated for 30 min at room temperature in binding/release buffer containing the indicated concentrations of $H_2O_2$ or 10 mM DTT. Ribosomes were precipitated by addition of trichloroacetic acid (TCA). Precipitated ribosomes were resuspended in 50 µl of alkylating buffer (200 mM IAA, 0.5 M Tris (pH 8.0), 5% glycerol, 100 mM NaCl, 2% SDS) and incubated for 1 h in the dark. Alkylated samples were precipitated by adding 1 ml of 10% TCA, resuspended in 100 µL 1.5 M Tris pH 8.0 and heat-denatured at 95°C for 10 min. Samples were cooled down to 37 °C and trypsin (Thermo Scientific; 1 µL of 0.5 µg/µL) digestion was performed in the presence of 1 mM $CaCl_2$ for 12 h. Samples were acidified with 1 vol of isopropanol/1% Trifluoroacetic acid (TFA) and desalted using styrenedivinylbenzene reverse-phase sulfonate (SDB-RPS) StageTips as described previously (*32*). Briefly, samples were loaded on a 200 µL StageTip containing two SDB-RPS disks and centrifuged at 1500 x g for 8 min. StageTips were washed three times with 200 µL of isopropanol 1% TFA at 1500 x g for 8 min, then eluted with 100 µL of 80% MeCN, 19% water, and 1% ammonia and dried. Peptides were resuspended in water with 0.1 % formic acid (FA) and analyzed using EASY-nLC 1200 nano-UHPLC coupled to Q Exactive HF-X Quadrupole-Orbitrap mass spectrometer (Thermo Scientific). The chromatography column consisted of a 45 cm long, 75 µm i.d. microcapillary capped by a 5 µm tip and packed with ReproSil-Pur 120 C18-AQ 2.4 µm beads (Dr. Maisch GmbH). LC solvents were 0.1 % FA in $H_2O$ (Buffer A) and 0.1 % FA in 90 % MeCN: 10 % $H_2O$ (Buffer B). Peptides were eluted into the mass spectrometer at a flow rate of 300 nL/min. over a 30 min long linear-gradient (5-35 % Buffer B) at 65 °C. Data was acquired in data-dependent mode (top-20, NCE 28, R = 15,000) after full MS scan (R = 60,000, m/z 300 – 1,650). Dynamic exclusion was set to 10 s, peptide match to prefer and isotope exclusion was enabled. The MS data were analyzed with MaxQuant (*33*) (V 2.0.3.0) and searched against the *Saccharomyces cerevisiae* proteome (Uniprot) and a common list of contaminants (included in MaxQuant). The first peptide search tolerance was set at 20 ppm, 10 ppm was used for the main peptide search and fragment

mass tolerance was set to 0.02 Da. The false discovery rate for peptides, proteins, and site identification was set to 1 %. The minimum peptide length was set to 6 amino acids and peptide re-quantification and "match between runs" was enabled. Following variable modifications were used: oxidation of methionine, protein N-terminal acetylation, carbamidomethylation of cysteine, and cysteine oxidation (+15.9949 Da), dioxidation (+31.9898 Da), and trioxidation (+47.9847 Da).

**Western analyses and Antibodies**

Western blots were scanned using the BioRad ChemiDoc MP Imaging System after applying luminescence substrates (Invitrogen) and quantified using its built-in image lab software (ver. 6.0.1). BTD-labeled protein imaging was performed using the Odyssey M Imaging System. The intensity of each band was analyzed after local background subtraction. For biotinylated BTD-probe detection, IRDye 800CW Streptavidin (LI-COR, 926-32230) and IRDye 689LT Goat anti-Rabbit IgG secondary antibody (LI-COR, 925-68021) was used. To detect TAP-tagged (Rps3) or HA-tagged proteins (Rps26 variants, Rpl10), anti-TEV cleavage site from Invitrogen (PA1-119) or anti-HA antibody from Abcam (ab18181) or Sigma (ab1603) were used, respectively. For Rps10 and Rps26 detection, antibodies were raised against a peptide by New England Peptide. Polyclonal antibodies were gifts from V. Panse (Tsr2/Rps26), M. Seedorf (Rps3), J. Warner (Rps2, Rpl3, Rpl32), B. Pertschy (Yar1) and KY Lo (Rpl23, Bcp1, Rpl43, Puf6, Loc1).

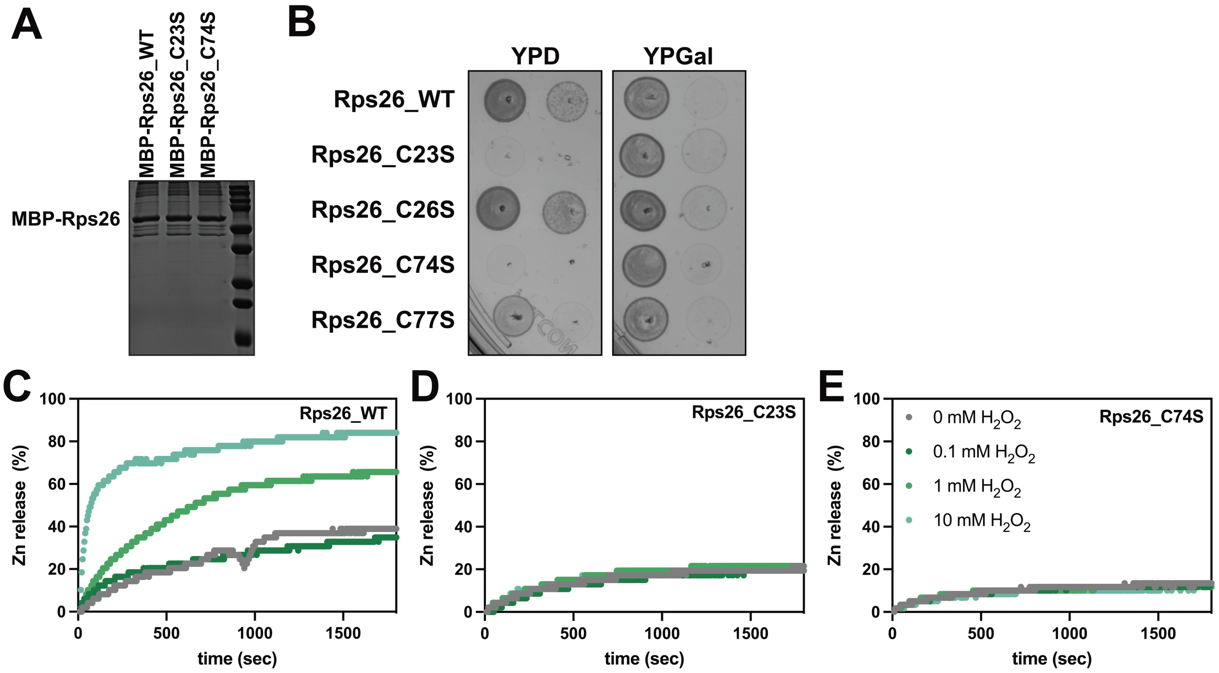


**Fig. S1.**

(A) Recombinant MBP-Rps26 variants used in Figure 1E and S1C-E. MBP-Rps26 variants were purified after overexpression in *E. coli*, and analyzed by SDS-PAGE. (B) Mutation of Rps26 cysteine residues leads to growth defects in yeast. YKK491 (Gal::Rps26A, *Δ*Rps26B) yeast cells containing plasmids encoding the indicated Rps26 variants were plated in 10 fold serial dilution either on glucose-containing media (to deplete endogenous Rps26) or on galactose media (to retain endogenous Rps26). (C-E) $H_2O_2$-dependent $Zn^{2+}$-release from Rps26. Release of $Zn^{2+}$ from 3.2 µM recombinant MBP-Rps26 (wt Rps26 or Rps26_C23S or Rps26_C74S) was measured by monitoring formation of the $Zn^{2+}$-PAR complex at 494 nm after treatment with 0, 0.1, 1 or 10 mM $H_2O_2$.

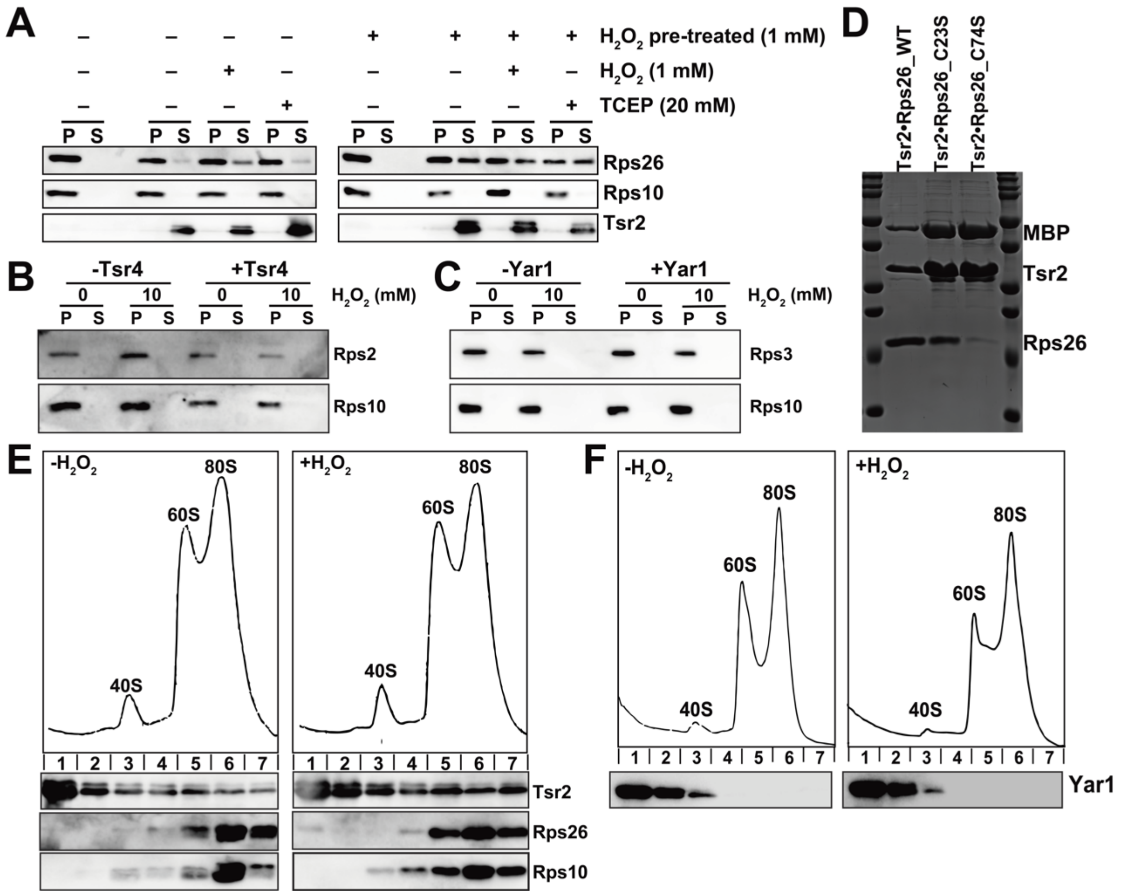


**Fig. S2.**

(A) Oxidation of Tsr2 is not required for the release of Rps26. 200 nM purified 40S subunits were incubated with or without 1 mM $H_2O_2$ for 30 min prior to pelleting. Pelleted ribosomes were resuspended in release buffer without (-) or with (+) $H_2O_2$ or TCEP as indicated, incubated with Tsr2 for 30 min. Western blot analysis was performed after a 2$^{nd}$ pelleting. S: supernatant; P: pellet. Rps2 (B) and Rps3 (C) are not released by their chaperones Tsr4 or Yar1. (D) Purified recombinant Rps26•Tsr2 (wt Rps26 or Rps26_C23 or Rps26_C74S) complex used in Figure 2C. (E) Recombinant Tsr2 binding to ribosomes with (right) or without (left) treatment with 1 mM of $H_2O_2$ for 30 min assayed in a 10-50% sucrose gradient followed by Western blot. (F) Recombinant Yar1 binding to ribosomes with (right) or without (left) treatment with 1 mM of $H_2O_2$ for 30 min assayed in a 10-50% sucrose gradient followed by Western blot.

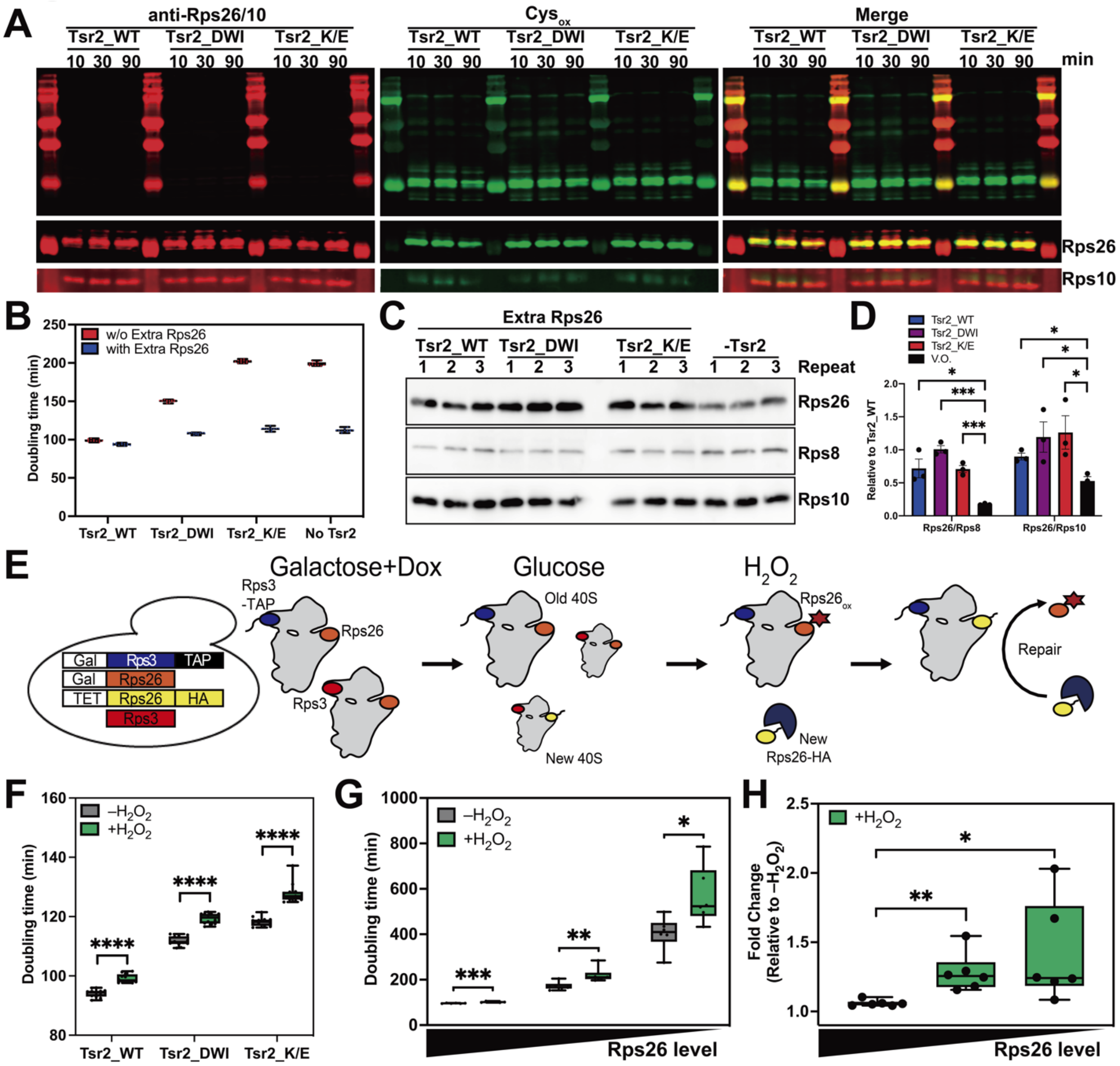


**Fig. S3.**

(A) Cells expressing wt Tsr2 or Tsr2_DWI or Tsr2_K/E were harvested in mid-log phase and pulsed for 5 min with 1 mM BTD and 1 mM $H_2O_2$. Cells were further cultured in 1L fresh media before harvesting at the indicated times. Ribosomes were purified from cell lysate by sucrose cushion and probed by Western blot. (B) Doubling time in glucose media of YKK1109 (Gal::Tsr2) cells expressing no Tsr2 or wt Tsr2 or Tsr2_DWI or Tsr2_K/E with or without overexpressing Rps26 from pKK3558. Data are the average of four biological replicates with three technical replicates. (C) Rps26 levels in ribosomes purified from YKK1109 (Gal::Tsr2) cells expressing wt Tsr2 or Tsr2_DWI or Tsr2_K/E and overexpressing Rps26 from pKK3558. Cells were grown overnight in glucose media. Rps8, Rps10 and Rps26 in purified ribosomes were determined by Western blot. (D) Quantification of 3 biological replicates in panel (C). (E) Scheme of pulse-chase experiments in Fig. 3F. To separate preexisting ribosomes from newly made ribosomes rely on a yeast strain with Rps3-TAP and Rps26 produced from a galactose-inducible/glucose-repressible (blue). By shifting this strain from galactose/dox to glucose, preexisting ribosomes are marked with the Rps3-TAP affinity purification handle and will be in the TAP-elution. Newly made

Rps26-HA (yellow) rely on a TET-repressible promoter to measure incorporation in Rps3-TAP tagged preexisting ribosomes. (F) Raw doubling times from Figure 3H. (G) Changes in doubling time upon addition 1 mM $H_2O_2$ in YKK491 cells (Gal:Rps26) with plasmid-encoded Rps26 produced in high (TEF-Rps26, left), moderate (TET-Rps26, mid) or low (TET-Rps26 + 25 ng/ml dox, right) concentrations. Data are the average of three biological replicates and two technical replicates. *$p < 0.05$, **$p < 0.01$, ***$p < 0.001$ by unpaired t-test. (H) Values normalized to untreated conditions from panel (G) (fold change = 1). *$p < 0.05$, **$p < 0.01$ by unpaired t-test.

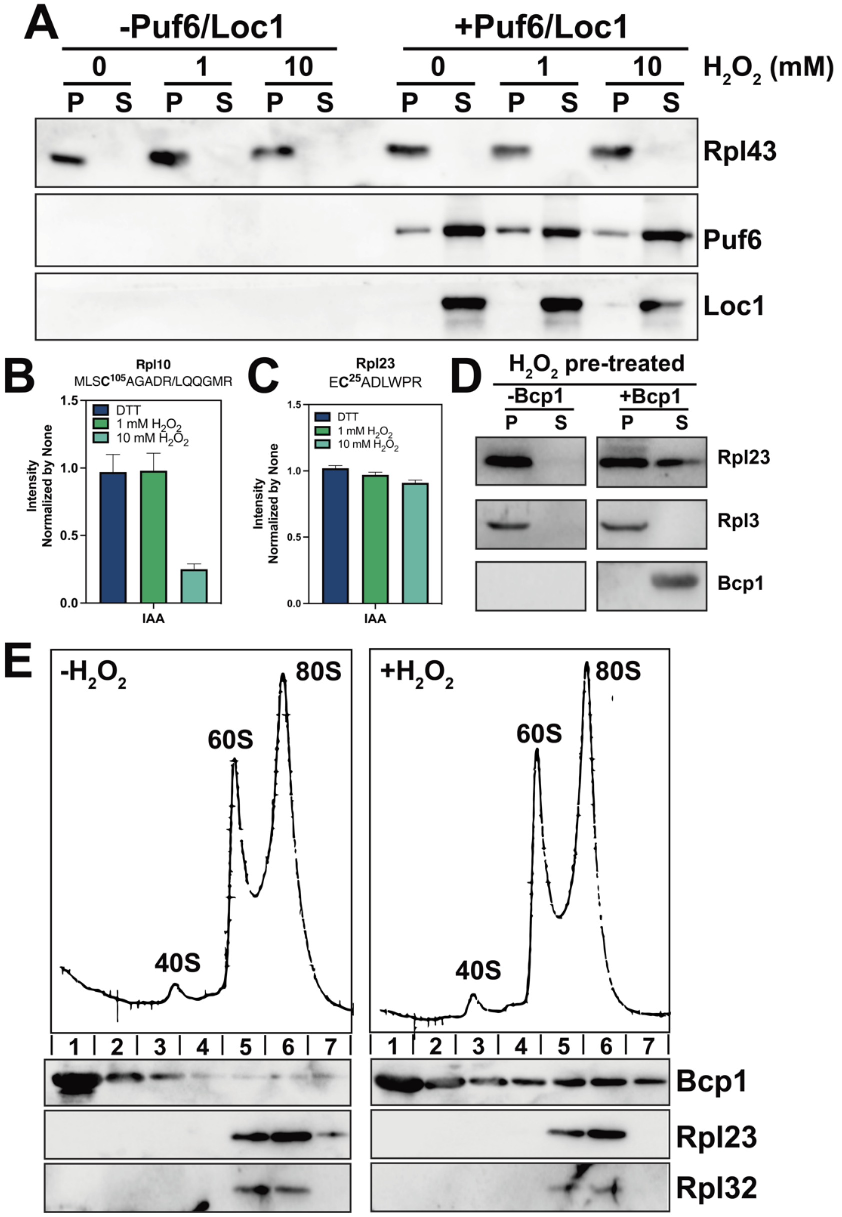


**Fig. S4.**

(A) Rpl43 is not released by chaperones Puf6/Loc1 *in vitro*. Mature 60S (200 nM) subunits purified from yeast were incubated with or without recombinant Puf6 and Loc1 (each 4 μM) under different $H_2O_2$ concentrations for 30 min prior to centrifugation. Release assay was performed in release buffer. S: supernatant; P: pellet. (B) Analysis of the Rpl10-derived peptide containing Cys105. (C) Analysis of the Rpl23-derived peptide containing Cys25. (D) Oxidation of Bcp1 is not required for the release of Rpl23. Purified 60S (200 nM) subunits were incubated with or without 10 mM $H_2O_2$ for 30 min prior to 1st step centrifugation. Pelleted ribosomes were resuspended in release buffer with or without Bcp1 and incubated for 30 min. Western blot analysis was performed after 2nd step ultracentrifugation pelleting experiments. S: supernatant; P: pellet. (E) Bcp1 interaction with ribosomes under oxidative stress. Western analysis of fractions from a 10-50% sucrose gradient of purified ribosomes from yeast lysates. 10 mM of $H_2O_2$ was treated (right) or untreated (left) for 30 min prior to sucrose gradients.

**Table S1.**
Yeast strains used in this work.

| Strain | Description | Background | Genotype | Reference |
|---|---|---|---|---|
| YKK200 | WT | BY4741 | *MATα his3Δ1 leu2Δ0 met15Δ0 ura3Δ0* | GE Dharmacon |
| YKK491 | Gal::Rps26 | BY4741 | *MATα NatMX6::pGAL1-Rps26A Rps26B::KanMX6 his3Δ1 leu2Δ0 met15Δ0 ura3Δ0* | (*21*) |
| YKK1109 | Gal::Tsr2 | BY4741 | *MATα NatMX6::pGAL1-Tsr2 his3Δ1 leu2Δ0 met15Δ0 ura3Δ0* | (*34*) |
| YKK1545 | Gal::Rpl10 | BY4741 | *MATα YLR075w::KANMX4 Leu2::pGAL1-Rpl10 his3Δ1 leu2Δ0 ura3Δ0* | (*35*) |

5

**Table S2.**
Plasmids used in this work.

| Plasmid | Description | Backbone | Reference |
|---|---|---|---|
| pKK3558 | TEF::Rps26A | pRS416 | (*21*) |
| pKK30832 | TEF::Rps26A_C23S | pRS416 | This work |
| pKK30831 | TEF::Rps26A_C26S | pRS416 | This work |
| pKK30832 | TEF::Rps26A_C74S | pRS416 | This work |
| pKK30831 | TEF::Rps26A_C77S | pRS416 | This work |
| pKK31045 | Gal::Rps3-TAP | pRS425 | This work |
| pKK30999 | TET:: Rps26-HA | pCM189 | This work |
| pKK30938 | TEF:: Rpl10-HA | pRS416 | This work |
| pKK30221 | TEF::Tsr2 | pRS415 | (*20*) |
| pKK30935 | TEF::Tsr2_K/E (K18E;K75E;K78E;K42E;K127E;K130;R140E;K142E;K143E;K145E;R146E) | pRS415 | (*20*) |
| pKK30570 | TEF::Tsr2_DWI | pRS415 | (*20*) |

**Data S1. (separate file)**
Peptides identified in *in vitro* oxidation of ribosomes.
